# Temperature affects recombination rate plasticity and meiotic success between thermotolerant and cold tolerant yeast species

**DOI:** 10.1101/2024.08.28.610152

**Authors:** Jessica McNeill, Nathan Brandt, Enrique J. Schwarzkopf, Mili Jimenez, Caiti Smukowski Heil

## Abstract

Meiosis is required for the formation of gametes in all sexually reproducing species and the process is well conserved across the tree of life. However, meiosis is sensitive to a variety of external factors, which can impact chromosome pairing, recombination, and fertility. For example, the optimal temperature for successful meiosis varies between species of plants and animals. This suggests that meiosis is temperature sensitive, and that natural selection may act on variation in meiotic success as organisms adapt to different environmental conditions. To understand how temperature alters the successful completion of meiosis, we utilized two species of the budding yeast *Saccharomyces* with different temperature preferences: thermotolerant *Saccharomyces cerevisiae* and cold tolerant *Saccharomyces uvarum*. We surveyed three metrics of meiosis: sporulation efficiency, spore viability, and recombination rate in multiple strains of each species. As per our predictions, the proportion of cells that complete meiosis and form spores is temperature sensitive, with thermotolerant *S. cerevisiae* having a higher temperature threshold for successful meiosis than cold tolerant *S. uvarum*. We confirmed previous observations that *S. cerevisiae* recombination rate varies between strains and across genomic regions, and add new results that *S. uvarum* has higher recombination rates than *S. cerevisiae*. We find that temperature significantly influences recombination rate plasticity in *S. cerevisiae* and *S. uvarum*, in agreement with studies in animals and plants. Overall, these results suggest that meiotic thermal sensitivity is associated with organismal thermal tolerance, and may even result in temporal reproductive isolation as populations diverge in thermal profiles.

## Introduction

The cell division process of meiosis is essential for all sexually reproducing species. While the core structures and processes are conserved across animals, plants, and fungi, genetic and environmental factors are known to mediate various aspects of meiosis including chromosome pairing and recombination (Lenormand et al., 2016; Wilkins & Holliday, 2009). Temperature is the most thoroughly studied of these environmental variables, with optimal temperature for meiosis varying across plant and animal species, and evidence of meiotic failure at high and low temperature extremes (Bomblies et al., 2015).

Variation in optimal meiotic temperature may in part reflect environmental sensitivity of meiotic structures. For example, the synaptonemal complex, which forms between homologous chromosomes during meiotic prophase I and facilitates chromosome pairing and recombination, shows defects and instability at high temperatures (Bayliss & Riley, 1972; Bilgir et al., 2013; Higgins et al., 2012; Loidl, 1989; Nebel & Hackett, 1961; Pao & Li, 1948; Zheng et al., 2014). Defects of the synaptonemal complex at high temperatures are likely linked to chromosome pairing failure, which results in improper segregation of chromosomes and reduced fertility (Elliott, 1955; Higgins et al., 2012; Loidl, 1989; Pao & Li, 1948; Yazawa et al., 2003). Genes encoding components of the synaptonemal complex and other meiotic proteins thus may be under selection as organisms experience different environmental conditions (Bomblies et al., 2015; Henderson & Bomblies, 2021; Morgan et al., 2017). Indeed, selection on meiotic genes has been identified in Drosophila, mammals, and Arabidopsis, though explicit links with temperature or the environment are not clear (Dapper & Payseur, 2019; Dumont, 2020; Samuk et al., 2019; Turner et al., 2008; Wright et al., 2015). Long standing laboratory studies in plants, animals, and yeast have documented recombination rate plasticity at different temperatures and support the connection between thermal sensitivity of meiotic processes and potential for meiotic evolution (Lim et al., 2008; Modliszewski et al., 2018; Morgan et al., 2017; Plough, 1917; Stevison et al., 2017; Winbush & Singh, 2021; Zhang et al., 2017). More recent studies in natural populations of Drosophila and Arabidopsis provide further correlational evidence, in which populations isolated from locations or seasons with different temperatures show differences in recombination rate (Dumont, 2020; Samuk et al., 2019; Weitz et al., 2021).

One lens through which we can better understand the link between temperature and the evolution of meiosis is to investigate how temperature alters meiotic phenotypes within and between related species with different thermal niches. The budding yeast *Saccharomyces* are an ideal system to investigate this question, as optimal growth temperature is a defining characteristic delineating species of the genus. Temperature appears to be an important factor in defining *Saccharomyces* species ranges and ecology (Langdon et al., 2020; Leducq et al., 2014; Robinson et al., 2016), and while little is known to impact prezygotic isolation between species, there is evidence that divergent thermal profiles may maintain distinct lineages in sympatry (Gonçalves et al., 2011; Sampaio & Gonçalves, 2008; Sweeney et al., 2004). The model system *Saccharomyces cerevisiae* is the most thermotolerant species, an apparent derived trait. It can grow in a wide range of temperatures, with a maximum temperature of 45.4°C (Salvadó et al., 2011). While known for its role in human associated fermentations like wine, beer, bread, and sake, it can also be isolated from natural environments including fruit, soil, and tree bark (Duan et al., 2018; Lee et al., 2022; Liti et al., 2009; Peris et al., 2022; Peter et al., 2018). Other species in the clade, like *Saccharomyces uvarum*, are more cold tolerant. *S. uvarum* has been isolated from cold-fermented beverages like wine and cider, and also is associated with tree bark (Almeida et al., 2014; Peris et al., 2022). Strains of *S. uvarum* have a maximum mitotic growth temperature of 36-38°C (Salvadó et al., 2011; Sampaio & Gonçalves, 2008), but some natural strain isolates appear more thermosensitive (Almeida et al., 2014).

Meiosis in *Saccharomyces* is facultative, and is induced when nitrogen and fermentable carbon sources are depleted, but non-fermentative carbon sources remain (Jambhekar & Amon, 2008; Zhao et al., 2018). Similar to gametogenesis in animals, the termination of the sexual cycle results in the formation of four haploid spores (this process is called sporulation). The ability and speed of sporulation is variable and heritable, and alleles at several genes have been identified to contribute to variation in this trait (De Chiara et al., 2022; Gerke et al., 2006; Tomar et al., 2013). Once sporulated, the spore wall is resistant to a variety of stressors and spores can remain in this dormant state for long periods of time. Sensing of glucose initiates the process of germination, and spores can resume asexual growth as haploids or mate to form diploids. Spore viability is intricately tied to successful chromosome pairing and recombination, with inviable spores often resulting from chromosome missegregation (Rogers et al., 2018).

While studies have identified genes and pathways implicated in mitotic temperature tolerance in *Saccharomyces* (AlZaben et al., 2021; Baker et al., 2019; Li et al., 2019; Weiss et al., 2018), meiotic thermal sensitivity is less understood. Common *S. cerevisiae* lab strains have variable sporulation at temperatures ranging from 15°C-34.5°C, with strain backgrounds S288C and W303 having reduced sporulation at higher and lower temperatures relative to 30°C (Elrod et al., 2009). A number of temperature sensitive alleles that cause meiotic arrest have been identified, but this has mostly focused on mutagenesis of lab strains and not identifying naturally segregating alleles (Byers & Goetsch, 1982; Davidow & Byers, 1984; Esposito & Esposito, 1969). As has been appreciated in a number of other organisms, temperature also changes the number and location of crossovers and non-crossover gene conversions in *S. cerevisiae* during meiosis (Cotton et al., 2009; Fan et al., 1995; Johnston & Mortimer, 1967; Zhang et al., 2017). Together, these studies suggest that meiosis is thermosensitive in *S. cerevisiae*, and that leveraging the strain and species diversity of the *Saccharomyces* clade may help untangle how selection acts on meiosis to shape adaptation to new temperatures, and how meiotic temperature tolerance contributes to reproductive isolation.

Given the important role temperature plays in delineating *Saccharomyces* species ranges and periods of activity, we hypothesize that there may be differences in optimal temperature for meiosis between species with different thermal preferences. Here, we utilized strains of thermotolerant *S. cerevisiae* and cold tolerant *S. uvarum* collected from a variety of ecological niches across the world (**Table S1**). We measured several meiotic phenotypes at a range of temperatures: the proportion of diploid cells that complete meiosis, the proportion of spores that are viable following meiosis, and recombination rate across multiple genomic intervals. We document variation within and between species in the successful completion of meiosis, and find that temperature affects recombination rate plasticity in both species.

## Results

### Mitotic growth recapitulates known thermal preferences in *S. cerevisiae* and *S. uvarum*

We selected 6 strains each of *S. cerevisiae* and *S. uvarum* that were isolated from different ecological niches and geographic locations (**Table S1**) (Almeida et al., 2014; Liti et al., 2009). We then crossed these *S. uvarum* and *S. cerevisiae* strains of interest to fluorescently-marked recombination tester strains with known lab strain backgrounds of CBS7001 and SK1, respectively, which we used in all subsequent experiments. To investigate the effect of temperature on the mitotic growth kinetics of these crosses, we measured the growth rate of one representative diploid from each set of *S. cerevisiae* and *S. uvarum* fluorescent strain crosses at three different temperatures (25°C, 30°C, 37°C) (**Figure S1, Tables S4-5**). Microbial growth rate is both genetically and environmentally labile, varying widely between strains and conditions. Generally, we recapitulate known thermal preferences for both species. *S. uvarum* exhibited negligible growth at 37°C for any strain except CBS7001, which had an average growth rate of 0.233 (SD 0.0755), and thus was excluded from further analysis. Temperature has a significant effect on growth rate in *S. uvarum*, with growth rate higher at 30°C than 25°C degrees (anova p<2.2×10-16), and in *S. cerevisiae*, with growth rate lower at 25°C and similar at 30°C and 37°C (Welch’s anova, p<1.6×10-16, Games-Howell test p<0.05). As expected, *S. cerevisiae* grows faster than *S. uvarum* at 25°C and 30°C (2-way anova, p<2.2×10-16 for temperature, species, and p<0.0016 for their interaction).

We interpret these results to be consistent with previous observations that *S. cerevisiae* has evolved to become more thermotolerant. While *S. cerevisiae* grows at a higher rate than *S. uvarum* at all temperatures surveyed, it is likely that *S. cerevisiae* would be outcompeted by more cold tolerant species like *S. uvarum* at colder temperatures (below 15°C). This is supported by results from competitive growth assays between sympatric isolates of *S. cerevisiae* and *S. kudriavzevii*, in which *S. cerevisiae* outcompetes *S. kudriavzevii* at 30°C but *S. kudriavzevii* outcompetes *S. cerevisiae* at 6°C (Sampaio & Gonçalves, 2008).

### High temperatures result in meiotic failure in cold tolerant *S. uvarum*

To identify how temperature affects meiosis in *S. cerevisiae* and *S. uvarum*, we first measured the proportion of diploid cells that complete meiosis and form spores (sporulation efficiency) (**Figure 1A, Tables S6-S7**). We induced meiosis at temperatures 4°, 10°, 15°, 25°, 30°, 37°, and 42°C (*S. cerevisiae* strains only) for 10 days (see Methods). There is extensive *S. cerevisiae* strain variation in ability to sporulate, and the length of time needed to sporulate, with several major loci known to contribute to variation (De Chiara et al., 2022; Gerke et al., 2006; Tomar et al., 2013). In accordance with previous research performed by Raffoux et al. (2018b), we deemed an incubation period of at least 10 days sufficient to capture the maximum sporulation efficiency of all *S. cerevisiae* and *S. uvarum* strains under study at each temperature. No strains from either species were able to produce appreciable (>3%) spores at 4°C, establishing this temperature as beyond the lower thermal boundary of successful sporulation for both *S. cerevisiae* and *S. uvarum.* Most *S. cerevisiae* strains were able to successfully sporulate within the range of 10°C to 37°C, with the exception of the YILC17_E5 cross, which exhibited a far narrower range of only 15°C to 30°C. Two *S. cerevisiae* strain crosses, SK1 and DBVPG6044, also proved capable of producing spores at an upper thermal extreme of 42°C. The SK1 cross exhibited the highest average sporulation efficiency of the species across all temperatures under survey (74.79% when averaged across all replicates, including 4°C; all replicates were included in this calculation to evaluate the comparative robusticity of strain sporulation across all thermal environments), and remained among *S. cerevisiae* strains with the highest average sporulation efficiency at each temperature. Such a high degree of consistent meiotic success in this cross is expected, as SK1 is a well-established *S. cerevisiae* lab strain that has been selected for use in studies of meiosis. With a truncated thermal range of sporulation compared to all other *S. cerevisiae* strains, the YILC17_E5 cross displayed the lowest average sporulation efficiency of the species across all temperatures (19.93% when averaged across all replicates, including 4°C). Even at temperatures where the YILC17_E5 cross was able to successfully produce spores, the cross consistently exhibited the lowest average sporulation efficiency of the species, reaching a maximum average of only 51.83% (SD=0.91) at 30°C.

**Figure 1.**
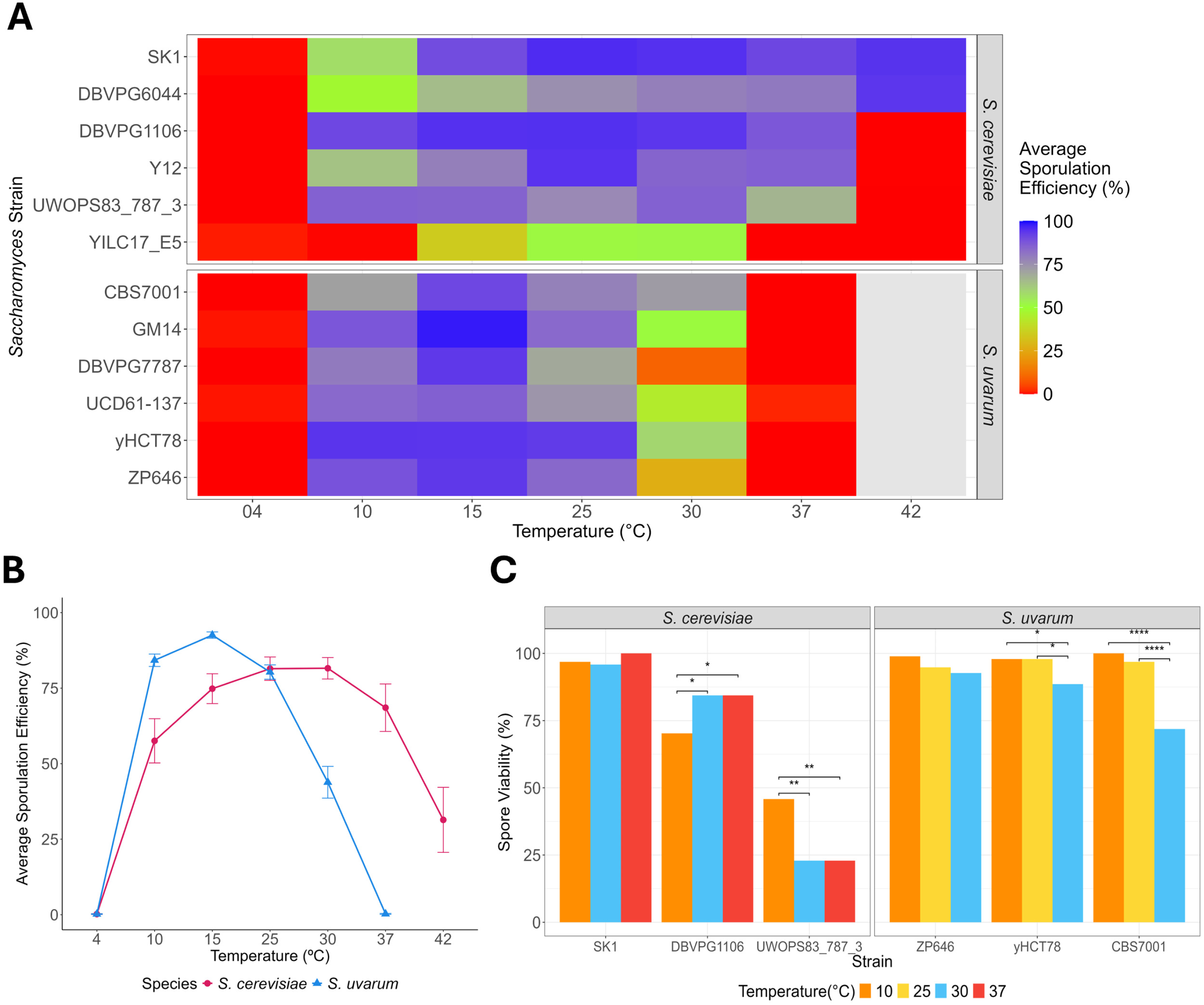
Sporulation efficiency and spore viability of *S*. *cerevisiae* and *S. uvarum* strains as crossed with a fluorescent tester of a lab strain background (SK1 and CBS7001, respectively). Diploid strains are labeled in reference to their non-fluorescent parent. **A.** Heatmap displaying average sporulation efficiency (%) of each strain calculated from 3 biological replicates sporulated at each temperature for a minimum of 10 days. No data was collected for *S. uvarum* strains at 42°C. **B.** Average sporulation efficiency (%) of each species at each measured temperature. Error bars indicate standard error across all species’ strain replicates. **C.** Spore viability (%) calculated from at least 21 meioses (84 spores) of 3 selected strains from each species. Temperatures were selected to correspond with the known thermal preference and observed lower and upper boundaries of successful sporulation within each species. Stars denote significance as revealed through a fisher exact test (assuming a 95% confidence interval), followed with a post-hoc pairwise Fisher’s exact test(p values corrected using Benjamini-Hochberg FDR method for at a 5% cut-off).

All *S. uvarum* strains were able to produce spores within the range of 10°C to 30°C; however, average sporulation efficiencies at 30°C dropped as low as 9.54% (DBVPG7787, SD=1.73). Unlike *S. cerevisiae,* no *S. uvarum* strains were able to produce appreciable spores at 37°C, establishing this temperature as above the upper thermal boundary of successful sporulation in *S. uvarum.* The yHCT78 cross displayed the highest average sporulation efficiency of the species across all temperatures (56.71% when averaged across all replicates, including 4° and 37°C; maximum 94.24%, SD=1.51 at 10°C). The lab strain CBS7001 cross reached a maximum average sporulation efficiency of 90.82% (SD=2.91) at 15°C, comparable to the maxima of most other *S. uvarum* crosses.

Because solid versus liquid media is known to influence sporulation efficiency, we also measured the sporulation efficiency of the lab strain crosses of both species (SK1 and CBS7001 for *S. cerevisiae* and *S. uvarum,* respectively) when cultured in liquid sporulation media. Average sporulation efficiencies for liquid *S. cerevisiae* SK1 cultures were higher than their solid media counterparts at 10°C and comparable at 15°C; however, these measures appeared lower in liquid media at higher temperatures, starting at 25°C (**Table S8**). A marked decline in sporulation efficiency appears most notable at 37°C, at which liquid sporulation cultures measured an average sporulation efficiency of only 5.77% (SD=2.79) in contrast to the average measure of 90.88% (SD=0.20) obtained on solid media. The *S. uvarum* CBS7001 strain had comparable average sporulation efficiencies to solid cultures at both 15° and 25°C. At 10°C, sporulation efficiency measures were both higher and less varied, with an average of 90.32% (SD=0.76) in liquid compared to 71.42% (SD=10.81) on solid media. However, like SK1, CBS7001 sporulation measures also starkly declined in liquid media near the upper limit of the species’ thermal range, measuring an average sporulation efficiency of only 5.77% (SD=2.64) in contrast to 78.36% (SD=8.72) on solid media. While sporulation efficiency does differ depending on media type, we find that the pattern of an increased thermal limit for sporulation in *S. cerevisiae* compared to *S. uvarum* holds true.

Overall, when we take an average sporulation efficiency of all strains from each species at each temperature, we see a parabolic effect of temperature on sporulation efficiency (**Figure 1B**). The shape of the parabola is dependent on species, with *S. cerevisiae* showing a peak in sporulation efficiency at 25°C and 30°C, while *S. uvarum* shows a peak at 15°C. We identify a significant effect of temperature (and temperature squared) by species interaction (**Table 1**), which supports our hypothesis that thermotolerant *S. cerevisiae* can produce spores at higher temperatures than cold tolerant *S. uvarum*.

Following our analysis of sporulation efficiency, we dissected tetrads from a subset of *S. cerevisiae* and *S. uvarum* strains to calculate the proportion of cells that survive meiosis and germinate (spore viability) at temperatures across our observed range of successful sporulation (**Figure 1C, Tables S9-S10**). Reduced spore viability is often the result of chromosome missegregation and aneuploidy (Rogers et al., 2018), which has also been linked to temperature sensitivity, though not yet in yeast (Henderson & Bomblies, 2021). We induced meiosis at 10°, 30°, and 37°C for *S. cerevisiae* strains, and 10°, 25°, 30°C for *S. uvarum -* temperature ranges that correspond with each species’ known thermal preference, flanked by temperatures approaching the observed lower and upper boundaries of successful sporulation within each species.

Analyses of *S. uvarum* strains show a consistent pattern of reduced spore viability with increasing temperature. While we detected no significant association between temperature and spore viability for the *S. uvarum* ZP646 cross (Fisher’s Exact, p=0.0915), we identified a significant association between these factors in the CBS7001 cross (Fisher’s Exact, p=1.11e-11) as well as the yHCT78 cross (Fisher’s Exact, p=5.31e-9). For both crosses, spore viability is lower at at 30°C than 10°C (Fisher’s Exact post-hoc FDR CBS7001: p-adj=0.5.31e-9; yHCT78: p-adj=0.0274), and 25°C (Fisher’s Exact post-hoc FDR CBS7001: p-adj=2.26e-6; yHCT78: p-adj=0.0274). This suggests that temperature impacts the spore viability of these *S. uvarum* strains at the upper boundary of their sporulation temperature range. In analyses of the *S. cerevisiae* strain subset, we detected no significant association between temperature and spore viability for the *S. cerevisiae* SK1 cross (Fisher’s Exact, p=0.1353); however, we did detect a significant association between these factors in the UWOPS83_787_3 cross (Fisher’s Exact, p=0.0005), as well as within the DBVPG1106 cross (Fisher’s Exact, p=0.0343). For both strains, this association is significant in comparisons between spore viability at 10°C and 30°C (Fisher’s Exact post-hoc FDR UWOPS83_787_3: p-adj=0.002; DBVPG1106: p-adj=0.0459) and at 10°C and 37°C (Fisher’s Exact post-hoc FDR UWOPS83_787_3: p-adj=0.002; DBVPG1106: p-adj=0.0459). Thus, temperature appears to have some impact on the spore viability of these strains at the lower boundary of their sporulation temperature range, but the nature of this impact appears to be strain dependent.

### Temperature affects recombination rate plasticity in *S. cerevisiae* and *S. uvarum*

Finally, we measured the effect of temperature on recombination rate plasticity. We utilized *S. cerevisiae* tester strains marked with fluorescent markers (Raffoux et al., 2018a), and created two *S. uvarum* fluorescent tester strains to correspond to syntenic intervals in *S. cerevisiae* for comparison. We crossed each *S. cerevisiae* and *S. uvarum* strain to the corresponding species fluorescent tester strains and induced meiosis for all crosses at temperatures 10°, 15°, 25°, 30°, and 37°C, and 42°C (*S. cerevisiae* only) for 10 days, as above. The temperature 4°C was excluded for both species since cells failed to sporulate at that temperature. After 10 days, sporulated cultures were enriched for spores and analyzed for recombination using flow cytometry. Altogether, we measured recombination rate for six strains of *S. cerevisiae* in 10 genomic intervals ( chromosomes I, VI, and XI) at six temperatures and five strains of *S. uvarum* in two genomic intervals (chromosomes VI and XI) at four temperatures (**Figure 2, Figures S2-S5, Tables S11-12**). We excluded several strain/interval/temperature combinations due to low sporulation or other technical issues.

**Figure 2.**
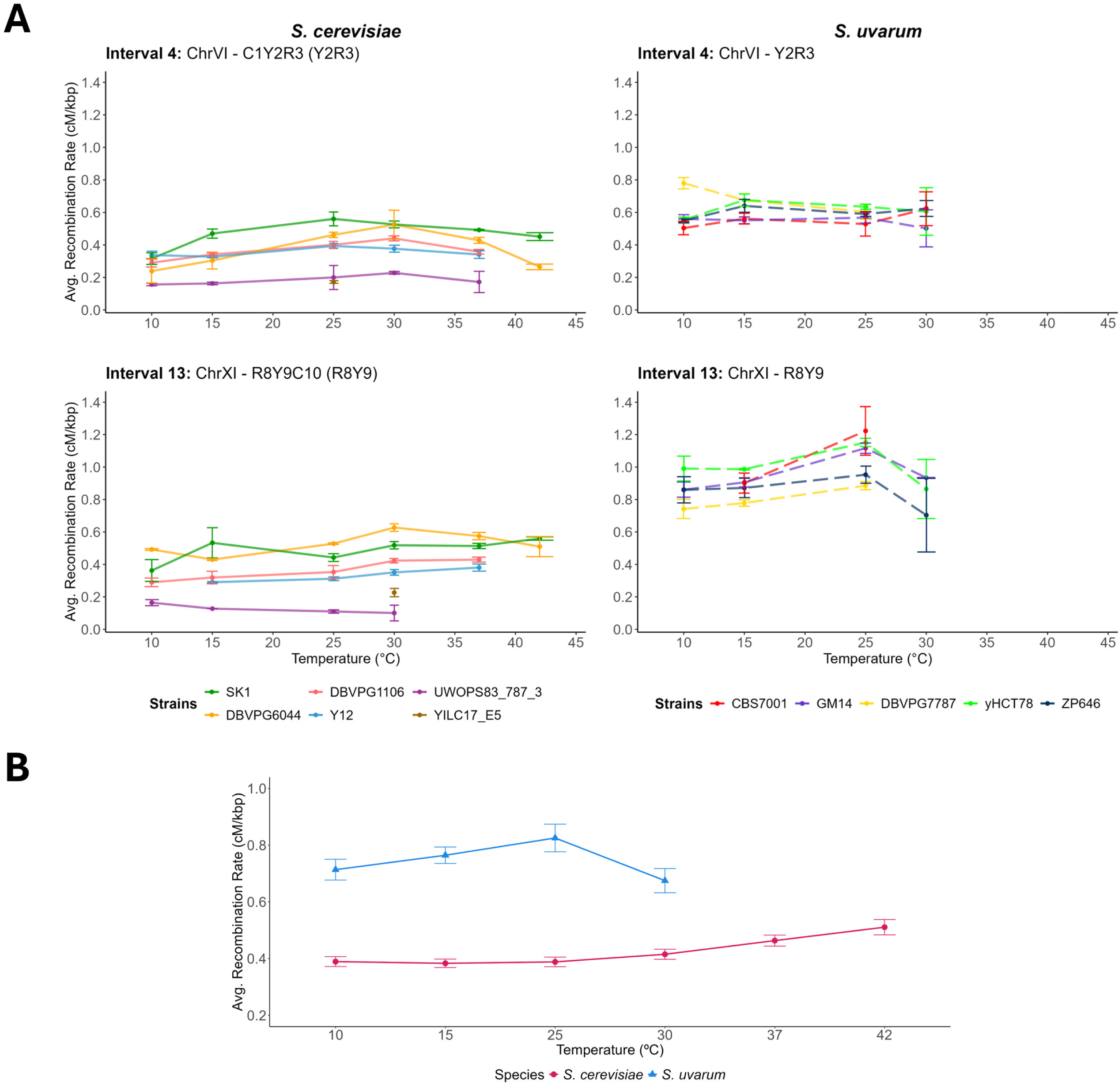
Average recombination rate (cM/kb) calculated for the syntenic intervals of 4 (on ChrVI) and 13 (on ChrXI) in *S. cerevisiae* and *S.uvarum* across all measured temperatures. *S. cerevisiae* recombination rate estimates were corrected for fluorescence extinction using a maximum likelihood model derived in Raffoux et al 2018a. *S. uvarum* recombination rate estimates were corrected for fluorescence extinction using a derivative of this script adjusted for only two fluorescent loci. No data was collected for *S. uvarum* strains at 42°C. In strain-level comparisons ***(A)***, strain name refers to the parent strain crossed with each fluorescent tester to produce a diploid for sporulation. Error bars indicate standard deviation above and below the mean, as calculated between biological replicates of each strain. In species-level comparisons, ***(B)*** error bars indicate standard error across all species’ strain replicates.

Previous work documented significant effects of strain and genomic region on *S. cerevisiae* recombination rates across measured intervals on chromosomes I, VI, and XI when sporulated at 30°C (Raffoux et al., 2018b). We first confirmed that we recapitulated *S. cerevisiae* recombination rates measured in that study at 30°C, and our results largely agree. In accordance with Raffoux et. al.’s findings, when we pooled and averaged *S. cerevisiae* data from across all intervals measured for each strain at 30°C (a maximum of 10 intervals), the homozygous SK1 cross retained the highest “global” recombination rate (0.58 cM/kb), followed closely by the DBVPG6044 hybrid (0.54 cM/kb), and more distantly by the DBVPG1106 and Y12 crosses (0.47 and 0.41 cM/kb, respectively). The UWOPS83_787_3 and YILC17_E5 crosses presented the lowest average global rates (0.22 and 0.18 cM/kb, respectively). This ordering remains relatively consistent across “global” recombination rates calculated at all other temperatures, with SK1 retaining the highest average value at all temperatures except 10 and 25°C, at which SK1 and DBVPG6044 rates are comparable (**Figures S6, S2-S5**).

Overall, we see a clear quadratic effect of temperature on recombination rate (**Figure 2B**, **Table 2**), consistent with studies in many plants and animals that temperature influences recombination rate plasticity. *S. uvarum* recombination rates peaks at 25°C, with lower recombination rates at higher and lower temperatures, whereas *S. cerevisiae* recombination rates increase with temperature, peaking at 42°C. However, we found no significant effect of the interaction of species and temperature or temperature squared, indicating that the effect of temperature on recombination rate is not significantly different between species. This may be influenced by our inability to measure recombination rates at higher temperatures in *S. uvarum* due to low sporulation rates. Interestingly, the temperatures with the highest sporulation efficiency and the temperatures with the highest recombination rates differ for both species, with optimal sporulation efficiency occurring at lower temperatures than highest recombination rates (**Figure 1B**, **Figure 2B**).

Average *S. uvarum* recombination rates are significantly higher than *S. cerevisiae* (**Figure 2**, **Table 2, Table S12**). While this only considers two genomic intervals, the high rate of *S. uvarum* recombination is consistent with our previous work finding high genome wide recombination rates in *S. uvarum* (Schwarzkopf et al., 2024). In comparisons between the two *S. uvarum* intervals using data averaged across all strains and temperatures, Interval 13 on ChrXI exhibited higher rates of recombination than Interval 4 on Chr VI (0.93 cM/kb, SD=0.15 compared to 0.59 cM/kb, SD=0.08). This disparity is observable across *S. uvarum* strain averages at each temperature, but is most apparent at 25°C, where the average recombination rate of Interval 13 (1.07 cM/kb, SD=0.14) is nearly double that of Interval 4 (0.58 cM/kb, SD=0.05). Strain yHCT78, isolated from tree bark at a vineyard in Missouri, USA, has the highest recombination rates averaged across all temperatures (Interval 4: 0.62 cM/kb, Interval 13: 1.01 cM/kb), whereas strains with the lowest recombination rates depend on the interval.

## Discussion

Sympatric species of *Saccharomyces* isolated from the same microhabitat have different thermal niches, as measured by their ability to grow mitotically under warm or cold temperatures (Gonçalves et al., 2011; Sampaio & Gonçalves, 2008; Sweeney et al., 2004). While most *Saccharomyces* species can grow at a wide range of temperatures, their optimal growth temperature and maximum growth temperature are clearly delineated, with *S. cerevisiae* outcompeting other species at warm temperatures and species like *S. kudriavzevii* and *S. uvarum* outcompeting at cold temperatures (typically at or below 12°C). Our study supports these observations, incorporating strains isolated from a variety of ecological niches, with *S. uvarum* exhibiting lower growth rates to *S. cerevisiae* at 25°C and 30°C, and unable to grow at 37°C.

In comparing meiotic phenotypes between thermotolerant and cold tolerant species, we predicted that as *S. cerevisiae* evolved to be increasingly thermotolerant, meiotic structures and processes would have increased thermal tolerance. Indeed, we do see an ability for *S. cerevisiae* to sporulate at higher temperatures than *S. uvarum*. Some studies have shown that selection for traits including geotaxis, pesticide resistance, parasite resistance, and temperature have resulted in increased recombination rates (Aggarwal et al., 2019; Kohl & Singh, 2018; Korol & Iliadi, 1994). We might extend this to predict that *S. cerevisiae* have higher recombination rates than *S. uvarum*, but we saw the opposite pattern. *S. uvarum* strains have higher recombination rates in the two syntenic intervals measured, which is in keeping with our recent work finding high genome wide recombination rates in *S. uvarum* (Schwarzkopf et al., 2024). However, the comparison between recombination rate evolution in *S. cerevisiae* and *S. uvarum* is complicated by their 20 million years of divergence, and recent domestication of Holarctic strains of *S. uvarum* used in this study (Almeida et al., 2014), which may also impact recombination rates (Bursell et al., 2024).

The effect of temperature on recombination rate plasticity is variable between different organisms, with a number of species displaying a U-shaped curve with the lowest recombination rate at ideal temperature conditions with elevated recombination rates at low and high temperatures (Bomblies et al., 2015; Henderson & Bomblies, 2021). We do not find a U-shaped pattern of recombination rate, instead we generally find an increase in recombination rate with temperature in both species (though *S. uvarum* recombination rate declines at the highest temperature able to be measured). No apparent patterns of chromosomal region or features like DSB, Rec8, sequence similarity, or H3K4me3 levels (Raffoux et al., 2018b) seem to be associated with which regions experience more plasticity. Under the hypothesis of fitness associated recombination rate plasticity, resistant genotypes may respond less than sensitive genotypes (Aggarwal et al., 2019; Rybnikov et al., 2017). For example, in a study that compared recombination rates of heat resistant and cold resistant tomato lines at different temperatures, the heat resistant line showed a more moderate increase in recombination rate than cold resistant lines at high temperature, and vice versa (Rybnikov et al., 2017). We did not find any significant effect of the interaction of species and temperature, and generally do not find support for fitness associated recombination rate plasticity.

Our meiotic phenotypes may also be impacted by other environmental variables, like sporulation on liquid vs. solid media and amount of sporulation time, which are known to affect sporulation efficiency and viability (Elrod et al., 2009). In previous work that documented recombination rate plasticity in *S. cerevisiae* due to temperature, recombination rates were measured using solid sporulation media with number of days sporulating varying by temperature (3-7 days) (Zhang et al., 2017). Zhang et al. (2017) did document an effect of genomic region on different recombination rates, with distance from the telomere and distance from the centromere having significant impacts on recombination rate plasticity. We do note that sporulation in liquid results in sporulation failure at a lower temperature than sporulation on solid media, however the pattern of effect of temperature on sporulation between species remains. By comparing meiotic phenotypes after 10 days, we are able to compare across genetic backgrounds without introducing additional variance due to effects on time to sporulation, although this surely is an additional component that would be interesting to pursue in future studies.

Our data further advance the idea that divergent thermal niches in *Saccharomyces* may be reinforced by meiotic failure at non-permissive temperatures, which provides a potential temporal reproductive isolating barrier between divergent populations and species. Prezygotic reproductive isolation barriers like temporal isolation between diverging populations may arise as a byproduct of local adaptation. For example, differences in flowering time as a result of selection due to varying soil moisture, altitude, and other climatic variables are a striking and highly repeated observation of temporal variation reducing gene flow (Lowry et al. 2008; MacNair and Gardner 1998; Vasek and Sauer 1971). While species in *Saccharomyces* have strong postzygotic isolation, prezygotic isolation is not well understood (Ono et al., 2020). Mating pheromones are conserved across all *Saccharomyces* (Rogers et al., 2015), and mate choice assays show little to no preference for conspecifics (Murphy et al., 2006; Naumov et al., 2000). Interspecific hybrids are frequently isolated from cold fermentations (Almeida et al., 2014; Dunn & Sherlock, 2008; Langdon et al., 2019), where they have a competitive advantage (Nikulin et al., 2018; Ortiz-Tovar et al., 2018). Almost all identified hybrids are between *S. cerevisiae* and a cold tolerant species, and it’s intriguing to speculate that conditions that favor interspecific mating are also related to temperature. Mating and germination timing do differ in sympatric species (Murphy & Zeyl, 2012), and it has been hypothesized that developmental timing and periods of activity may differ over the course of day or year (Sampaio & Gonçalves, 2008). Our results support this hypothesis, in which cold tolerant species would be unable to produce viable spores at certain times of the year.

Because temperature has such a strong influence on a wide variety of cellular processes and organismal behavior and survival, it has been studied extensively in a wide array of species. Organismal thermal tolerance across a range of temperatures is commonly used to measure and estimate range boundaries and predict response to climate change (Wooliver et al., 2022). However, many of these studies focus on parameters of growth that are not related to the sexual cycle, and an increasing number of studies support that an organism’s “thermal fertility limit” (TFL) is lower than its’ “critical thermal limit” (CTL) (Iossa, 2019; van Heerwaarden & Sgrò, 2021; Walsh et al., 2019). Our results add to these data, supporting that parental temperature growth maximums may overestimate an organism’s thermal niche, as meiotic failure occurs at a temperature below that of which it can sustain growth. We thus support recent work calling for measuring organisms’ thermal fertility limit to better understand population persistence in the face of environmental change (Walsh et al., 2019).

Finally, while meiosis has long been recognized to be sensitive to temperature, thermal sensitivity is rarely measured within species or between closely related species. Our documented variation of meiotic thermal sensitivity within and between species sets up a powerful system to test how meiotic thermal sensitivity can evolve. As the climate changes, this is likely to be a key parameter in determining species resiliency and long term survival.

## Materials and Methods

### Strains

A list of yeast strains is included in **Table S1**. Selected *S. cerevisiae* strains are part of the SGRP collection (Liti et al., 2009). The *S. uvarum* strains were obtained from the Portuguese Yeast Culture Center and courtesy of Chris Hittinger (Almeida et al., 2014). *S. uvarum* strains had their *HO* locus replaced with a kanMX or hygMX marker using a modified version of the high-efficiency yeast transformation using the LiAc/SS carrier DNA/PEG method with a heat shock temperature of 37°C for 45 minutes. The markers were amplified from plasmids pFA6a-TEF2Pr-dTomato-ADH1-Primer-KANMX6 and pUC-HygMX using primers (CSH239, GGTGGAAAACCACGAAAAGTTAGAACTACGTTCAGGCAAAgacatggaggcccagaatac; CSH240, TACTACTGGCAAATACACTAGCTTCTGGGCGAACTACAAGGTGGAAAACCACGAAAAGT; CSH241, GTGACCGTATTGGTACTTTTTTTGTTACCTGTTTTAGTAGcagtatagcgaccagcattc; CSH242, AAAACTTTTTGTTTGCATTCAATTATATCGATCAATGGAGTGACCGTATTGGTACTTTT) for multiplex PCR that introduced 60 bp of homology outside the *HO* ORF. Single colonies were selected from the transformation plates and sporulated and dissected as per methods above. The plates were incubated at room temperature for 2 days and then replica plated to test for proper segregation of antibiotic marker and mating type within individual tetrads.

### Measuring recombination rate

*S. cerevisiae* tester strains for measuring recombination rate were obtained courtesy of Matthieu Falque (Raffoux et al., 2018a, 2018b). Briefly, this set consists of an array of SK1 haploid strains (**Table S1**) built to contain three fluorophores spaced approximately 30 cM apart within regions of one of three chromosomes (chrI, chrVI, or chrXI, **Table S2**). Six *S. cerevisiae* strains that showed variation in recombination rate in previous work (Raffoux et al., 2018b) were selected from the SGRP collection (**Table S1**). Haploids of these six strains were then crossed via micromanipulation to fluorescent tester strains, and resulting diploids were confirmed using a halo assay. Following Raffoux et al, diploid strains were cultured overnight in 5mL of liquid YPD (1% yeast extract, 2% peptone, 2% dextrose) at 30°C. The following morning, each culture was transferred to a 15mL conical tube and centrifuged at 2000 rpm for 2 minutes. Cell pellets were resuspended in 450uL ddH_2_O, and 150uL of each mixture was spread onto solid SPM plates (1% Potassium Acetate, 0.1% yeast extract, 2% agar, 0.05% dextrose) in triplicate. These plates were then incubated at 4, 10, 15, 25, 30, 37, and 42°C for 10 days. All samples were then prepped for spore enrichment within the subsequent 3 days. To eliminate vegetative cells and enrich spores for recombination rate analysis, a methodology established by Raffoux et. al. was adapted with slight alteration. Following confirmation of sporulation by visual inspection, a quarter to a half of the lawn of each successfully-sporulated plate was harvested with a bent pipette tip and suspended in a 1.5mL tube containing 750uL of ddH_2_O with 5mg/mL 20T Zymolyase, after which 100uL of glass beads were added. To disrupt tetrads, this tube was vortexed for 1 minute at a frequency of 3000 rpm using a Disruptor Genie, incubated for 60 minutes at 30°C, then subjected to another 3000rpm vortex for 1 minute. The liquid portion of each mixture was then transferred into a new 1.5mL tube and centrifuged for 5 min at 4500rpm.

The resulting pellet was resuspended in 200uL of ddH_2_O, and this suspension was then discarded, as it contained primarily vegetative cells. The spores left adhering to the tube plastic were stripped by adding a solution of 600uL ddH_2_O with 0.01% NONIDET NP40 and vortexing for 30 seconds. Upon analysis on the flow cytometer, this concentrated spore suspension was diluted with up to 1mL of 1x PBS if necessary. Each culture was run on a Thermo Fisher Attune flow cytometer (University of North Carolina Flow Cytometry Core). To capture a population of spores, data were gated based on forward scatter and side scatter, then subsequently gated for single cells. All data yielding less than 2000 events within this single cell gate were discarded prior to further analysis.To confirm that the population of spores being captured was devoid of vegetative cells, mixed populations of vegetative and fluorescent spores, as well as vegetative cells, were analyzed as per Raffoux et al (2018a). Our spore enrichment protocol yielded around 1% carryover of vegetative cells. Distinct fluorescent populations were gated further to extract numeric counts of all possible genotypes within the total spore population; in accordance with the protocol derived in Raffoux et al 2018a, these counts could be input into a maximum likelihood model to estimate recombination rate while controlling for fluorescence extinction (Supplementary file “Text S1” from Raffoux et al., 2018a for *S. cerevisiae* testers with 3 fluorescent loci). If distinct fluorescent populations could not be resolved for a sample following single-cell gating, the data were discarded. An attempt was made to analyze all strain-by-temperature combinations, but insufficient spores and poorly resolved fluorescent populations were produced for certain strain backgrounds and temperatures **(Tables S6-7, S11-12**). Analysis of flow data was done using FloJo v10.10.0 (BD BioSciences).

To measure recombination rate in *S. uvarum*, two tester strains with two fluorophores on regions of either chrVI or chrXI were constructed using CBS7001 as a strain background (**Table S1, S2**). These intervals were designed to be homologous and syntenic to intervals in *S. cerevisiae*. Multiple attempts were made to create a third tester strain with syntenic intervals on chrI, but a fluorescent marker was unable to be integrated at locus 3. The TDH3prom-mCherry-KanMX construct was PCR-amplified from the I - R2C3Y4 strain of the *S. cerevisiae* fluorescent strain collection detailed above. Approximately 40 bp of locus homology was then added to either side of the construct through a second round of PCR using purified product from the first round of construct amplification as template. To bypass regions of shared sequence flanking both the TDH3prom-YFP-NatMX and TDH3prom-yECerulean-NatMX constructs, the *S. cerevisiae* I - R2C3Y4 strain was then crossed with a non-fluorescent SK1 strain, sporulated, and dissected to obtain a haploid retaining only the I-Y4 construct. The TDH3prom-YFP-NatMX construct was PCR-amplified from this I-Y4-only strain. For each fluor on each chromosome, the *S. uvarum* strains were transformed as above, following the same protocol as was used for the *ho* locus. Correct integration of each fluor was confirmed via PCR and flow cytometry. Primers used to amplify the constructs, add homology for transformation, and confirm correct locus integration post-transformation are listed in **Table S3**. mCherry and YFP strains were crossed and sporulated, and haploids of each mating type were confirmed to have both mCherry and YFP via PCR and flow cytometry. These haploids were then mated via micromanipulation to 6 *S. uvarum* strains of interest, and checked for diploidy using a halo assay. Diploid strains were cultured overnight in 5mL of liquid YPD at 25°C, and the protocol as above was used to plate strains onto triplicate solid SPM plates. Plates were then incubated at 4, 10, 15, 25, 30 and 37°C for 10 days. The same spore enrichment and flow cytometry protocol was followed as above for *S. cerevisiae*, accommodating for only two fluorescent markers per S. *uvarum* strain rather than the three present in the *S. cerevisiae* tester strains. An analogous maximum likelihood method was derived from the original Raffoux et al 2018a script to estimate recombination rate while controlling for fluorescence extinction at only two fluorescent loci (**Text S2**).

### Measuring growth rate

Each strain was grown in YPD overnight at room temperature. The following day, the culture was diluted to have a starting optical density (OD) of 0.1 or less and inoculated in 96 well plates at a volume of 200 ul. Growth curves were calculated for each *S. cerevisiae* and *S. uvarum* strain at temperatures 25°C, 30°C, and 37°C using a BioTek Epoch 96-well plate reader that measures the OD at 600 nm. Each strain was measured in 6 replicates (2 technical replicates for each of 3 biological replicates) in different wells of the plate for each run to account for plate positioning effects. OD readings were collected every 15 minutes over a 48-hour period with continuous dual orbital shaking. Growth was analyzed using the R package “growthcurver” to calculate growth rate (r) and area under the empirical curve (auc_e) (Sprouffske & Wagner, 2016). The sigma values are a measure of how well the model represents the growth curves and were used to identify potential outliers. Each potential outlier was then individually inspected to determine if the data point needed to be omitted.

### Sporulation efficiency

One representative diploid was chosen from each set of *S. cerevisiae* crosses and each set of *S. uvarum* strain crosses to be measured for sporulation efficiency at 4, 10, 15, 25, 30, and 37°C. *S. cerevisiae* representatives were subjected to an additional measure at 42°C. Strains were sporulated in triplicate following the same protocol as was used for the measure of recombination rate. After a 10-day incubation period, all measures were taken within the subsequent 3 days. A quarter of the lawn of each plate was harvested using a bent pipette tip and suspended in 1mL ddH_2_O. 10uL of this suspension was then placed onto a hemocytometer and investigated under a light microscope. If cell concentration remained too dense to facilitate clear counting at this stage, additional dilutions were performed. For each sample, two counts were taken of the number of spores with visible asci (a category including monads, dyads, and tetrads) within an observed population of at least 200 cells. The proportions resulting from these two counts were then averaged to control for any variation in spore calls between the two counts. Sporulation efficiency was then calculated from these averages as the percentage of spores with visible asci within the total population observed for each sample.

### Liquid sporulation

From the diploid strain subset utilized for the measure of sporulation efficiency on solid media, one representative strain was chosen from each species to be measured for comparative sporulation efficiency in liquid SPM media at 10, 15, 25, and 30°C. The *S. cerevisiae* representative was subjected to an additional measure at 37°C. From the *S.uvarum* strain set, a fluorescent cross with the lab strain CBS7001 was chosen. From the *S.cerevisiae* strain set, a fluorescent cross with the lab strain SK1 was chosen. Strains were sporulated using a protocol derived and modified from (Trainor et al., 2021). *S. uvarum* and *S.cerevisiae* strains were first streaked out onto YPD plates (1% yeast extract, 2% peptone, 2% dextrose, 2% agar) and grown for approximately 48-72 hours at 25° C and 30° C, respectively. A single colony was selected from each plate, moved to 5 mL liquid YPA (1% yeast extract, 2% peptone, 2% potassium acetate), and placed in a spinner to grow overnight at these same respective temperatures. This process was performed in triplicate to include 3 biological replicates for each cross at each temperature. The following day, a measure of OD600 was taken for each culture. Approximately 6 × 10^7^ cells were removed from each culture, spun down (∼2500 rpm for 1 minute), and washed with sterile ddH20. Cells were then resuspended in 1mL liquid SPM, and left to sporulate with agitation at a designated temperature for 10 days. After this 10-day incubation, 100 µl of each sporulated sample was diluted in 1 mL of sterile water, vortexed, and sonicated thoroughly. Cells were counted and percent sporulated cells calculated in the same method as described above.

### Spore viability

As above, strains were sporulated for 10 days at temperatures 10°C, 25°C, and 30°C (*S. uvarum*) or 10°C, 30°C, and 37°C (*S. cerevisiae*), and successful sporulation was checked under a light microscope. Once confirmed, a minute patch of cells from the lawn of the solid SPM plate was harvested with a bent pipette tip, and suspended in 50uL of ddH_2_O. This suspension was then centrifuged at 2000 rpm for 1 minute, and the resulting pellet was resuspended in a mixture of 25uL ddH_2_O and 2uL of 5mg/mL 20T Zymolyase (the equivalent of 1U). The tube was then incubated in a 37°C water bath for 10 minutes, immediately after which 500uL of ddH_2_O was added to halt the Zymolyase reaction. Tetrads were dissected on YPD plates using a Singer SporePlay+ microscope (Singer Instruments). While many sporulated cultures had a mix of monads, dyads, and tetrads, only complete tetrads were dissected. *S. cerevisiae* spores were grown for 2 to 4 days at 30°C, while *S. uvarum* spores were grown for an analogous time at room temperature. Spore viability was calculated as the number of spores that successfully returned to a vegetative state divided by the total number of spores dissected. A minimum number of 21 meioses (84 spores) were analyzed per temperature per strain; however, all but one sample successfully yielded 24 meioses (96 spores) for analysis.

## Data analysis and visualization

Statistical analyses were done using R packages “dplyr,” “FSA,” “car,” and “multcomp” (Fox et al., 2021; Hothorn et al., 2021; Ogle et al., 2021; Wickham et al., 2021). All plots were made using the R program ggplot2 (Wickham, 2016).

## Data and resource availability

*S. cerevisiae* fluorescent tester strains are courtesy of Matthieu Falque, *S. uvarum* strains are courtesy of the Portuguese Yeast Culture Collection (PYCC) and Chris Hittinger. *S. uvarum* fluorescent tester strains are available from our lab upon request. Raw data generated and analyzed in this paper are included in the supplementary tables.

## Acknowledgements

We thank Musa Malik and Adam Greer for collecting preliminary data on sporulation efficiency. Carrie Olson-Manning and Michael Law provided helpful comments on drafts of this manuscript. Thanks to Mathieu Falque, Chris Hittinger, and the Portuguese Yeast Culture Collection for sharing *S. cerevisiae* and *S. uvarum* strains. This work was supported by NIH R35 GM142849 to C.S.H.

## Author contributions

NB assisted in strain construction; JM performed experiments and collected data; MJ and CSH collected data; JM and EJS conducted data analysis; CSH supervised research; JM and CSH wrote the manuscript; all authors contributed to editing.

---

|  | Estimate | Std. Error | z value | Pr(>\|z\|) |
| --- | --- | --- | --- | --- |
| (Intercept) | -5.0323 | 0.7752 | -6.4915 | 0.00E+00 |
| Temperature | 0.574 | 0.081 | 7.0844 | 0.00E+00 |
| l(Temperature^2) | -0.0113 | 0.0017 | -6.7918 | 0.00E+00 |
| Temperature: Species S. uvarum | 0.274 | 0.0755 | 3.6308 | 3.00E-04 |
| l(Temperature^2): Species S. uvarum | -0.0117 | 0.0027 | -4.4002 | 0.00E+00 |

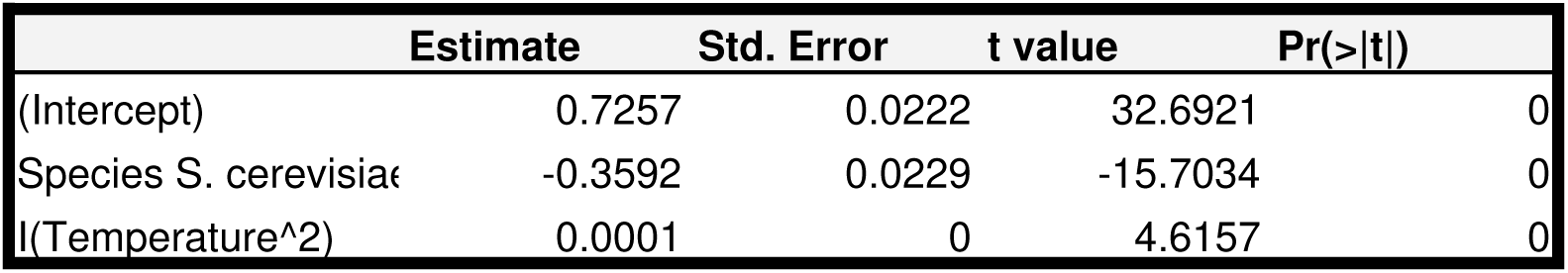

