## Supplemental Information for "Temperature affects recombination rate plasticity and meiotic success between thermotolerant and cold tolerant yeast species"

**Figure S1**

Growth rate (r) as calculated for the mitotic growth of each strain in liquid YPD. All strains were diploid crosses between a strain of interest (total 6 for each species) and a recombinant fluorescent tester strain of a lab strain background (SK1 for *S. cerevisiae*, CBS7001 for *S.uvarum*). Diploid strains are labeled in reference to their non-fluorescent parent. At 37°C, *S.uvarum* strains failed to produce a measurable growth curve; thus, these data have been omitted from those plotted.
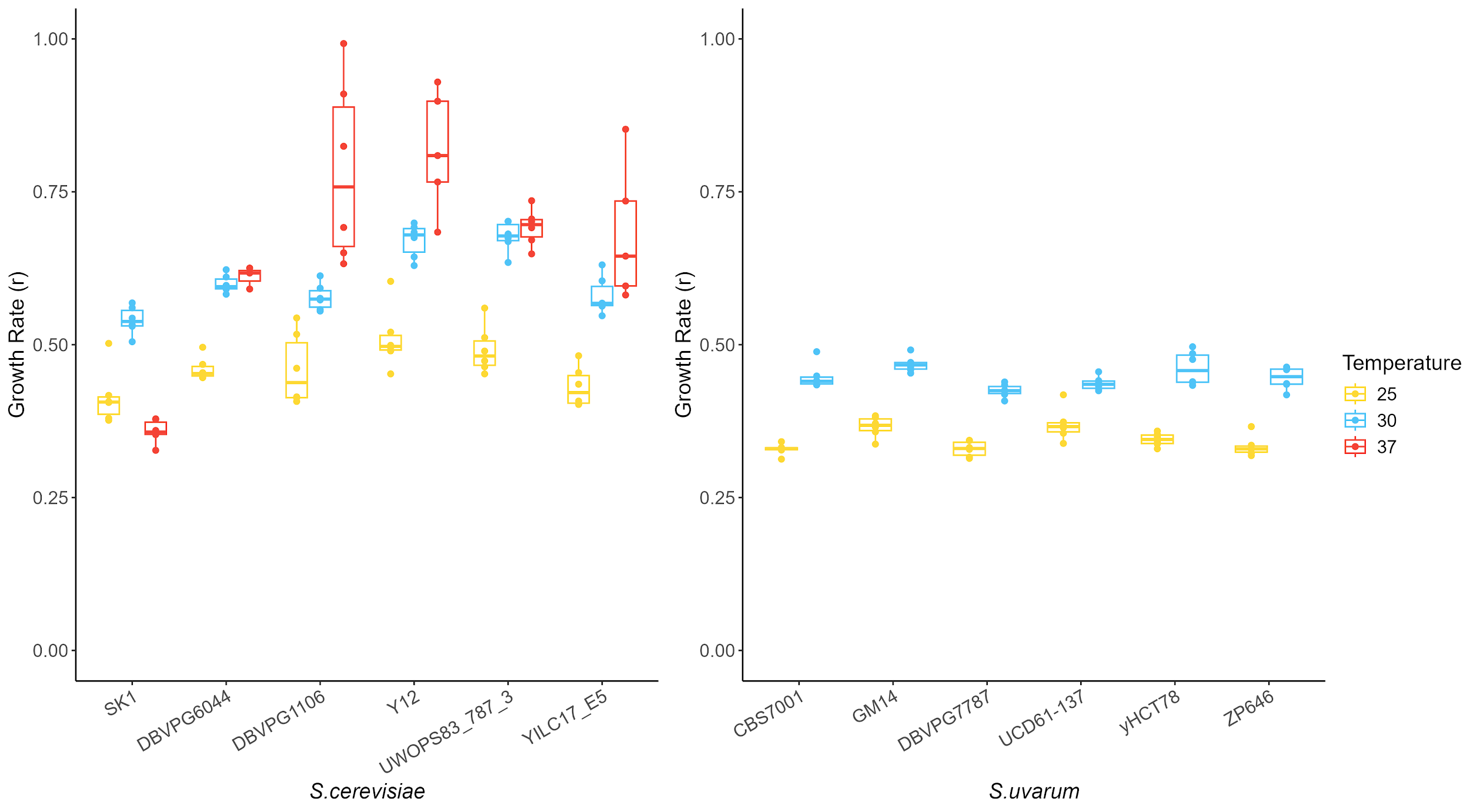

**Figure S2**

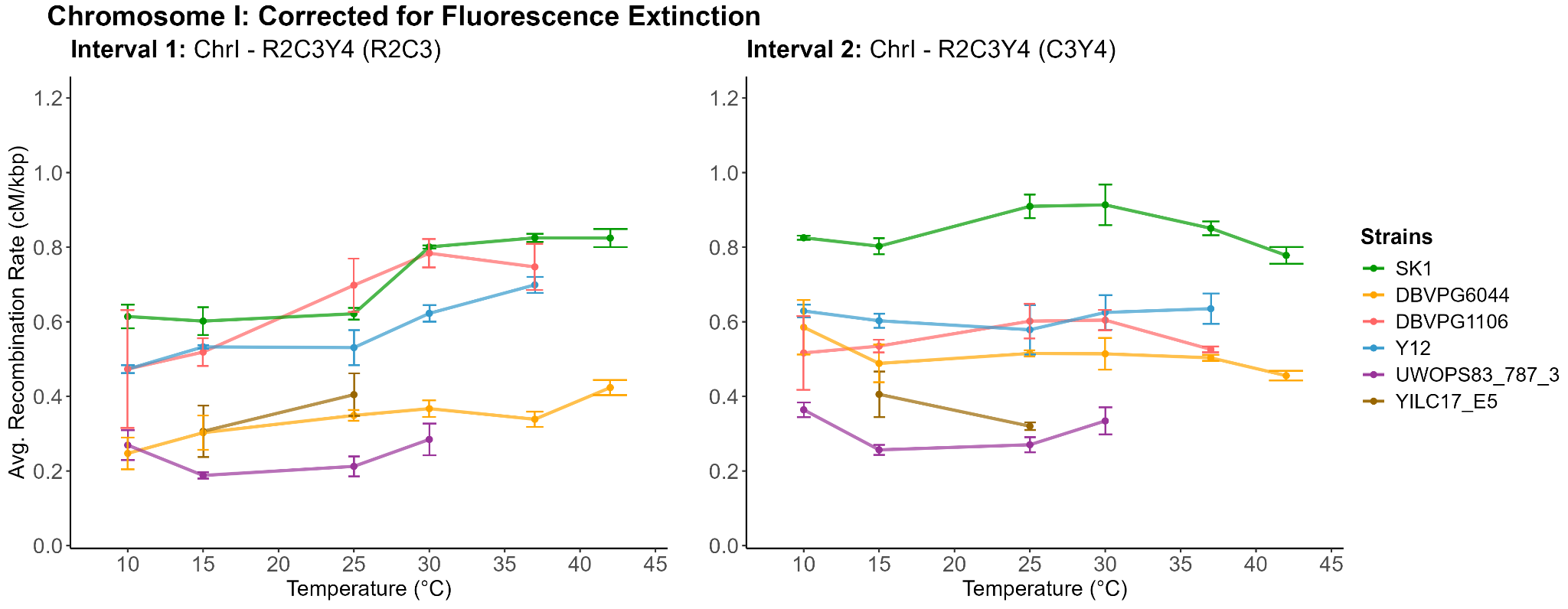

Average recombination rate (in cM/kbp) as calculated for intervals 1 and 2 from sporulated crosses with the ChrI-R2C3Y4 fluorescent tester strain at all viable temperatures. Strain name refers to the parent strain crossed with this fluorescent tester to produce a hybrid diploid. Recombination rate estimates were corrected for fluorescence extinction using a maximum likelihood model derived in Raffoux et al 2018a. Error bars indicate standard deviation above and below the mean, as calculated between biological replicates.

**Figure S3**

Average recombination rate (in cM/kbp) as calculated for intervals 3, 4 ,5 and 6 from sporulated crosses with the ChrVI-C1Y2R3 fluorescent tester strain (3 and 4) and the ChrVI-R3Y4C5 fluorescent tester strain (5 and 6) at all viable temperatures. Strain name refers to the parent strain crossed with each fluorescent tester to produce a hybrid diploid. Recombination rate estimates were corrected for fluorescence extinction using a maximum likelihood model derived in Raffoux et al 2018a. Error bars indicate standard deviation above and below the mean, as calculated between biological replicates.
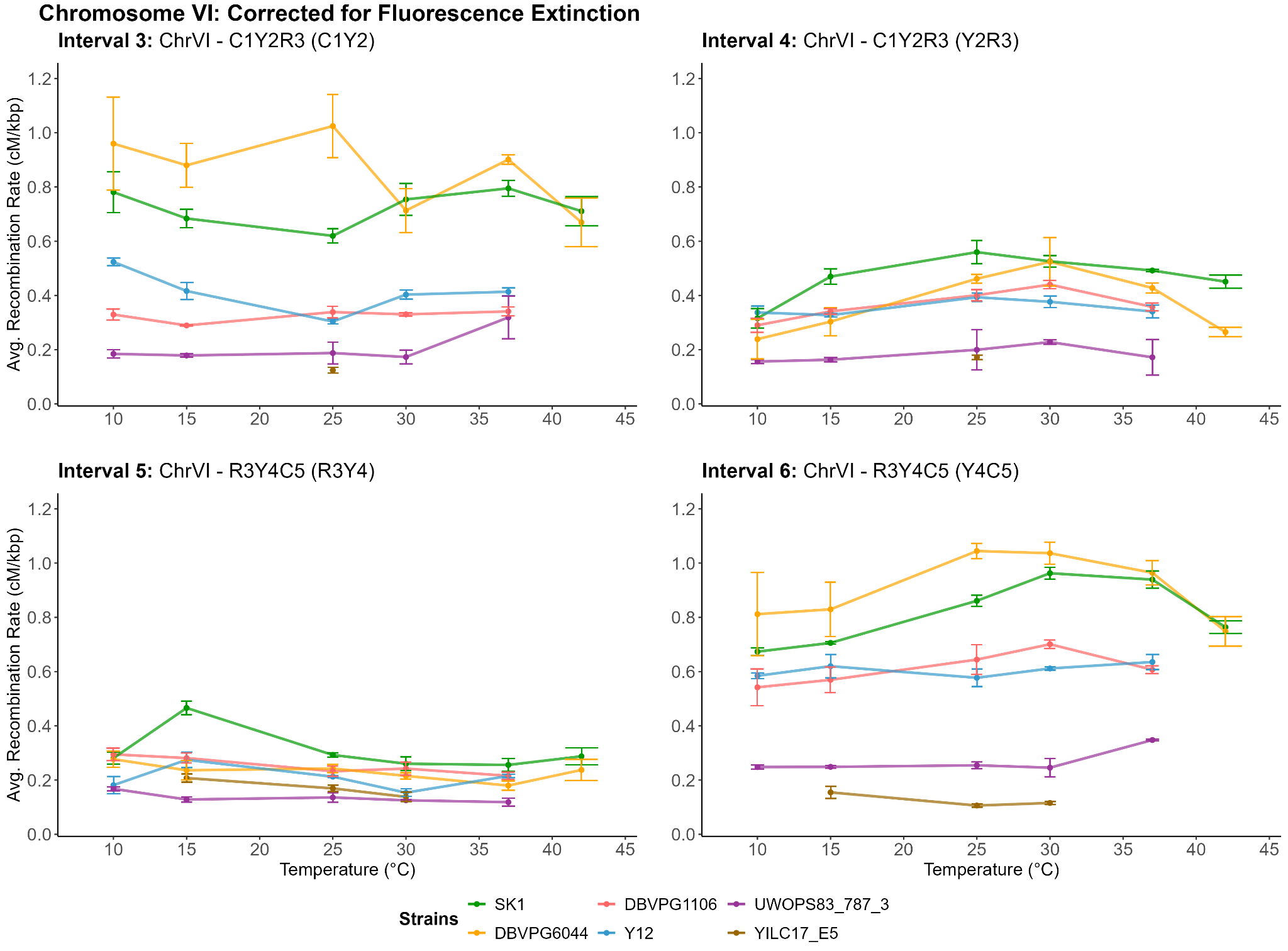

**Figure S4**

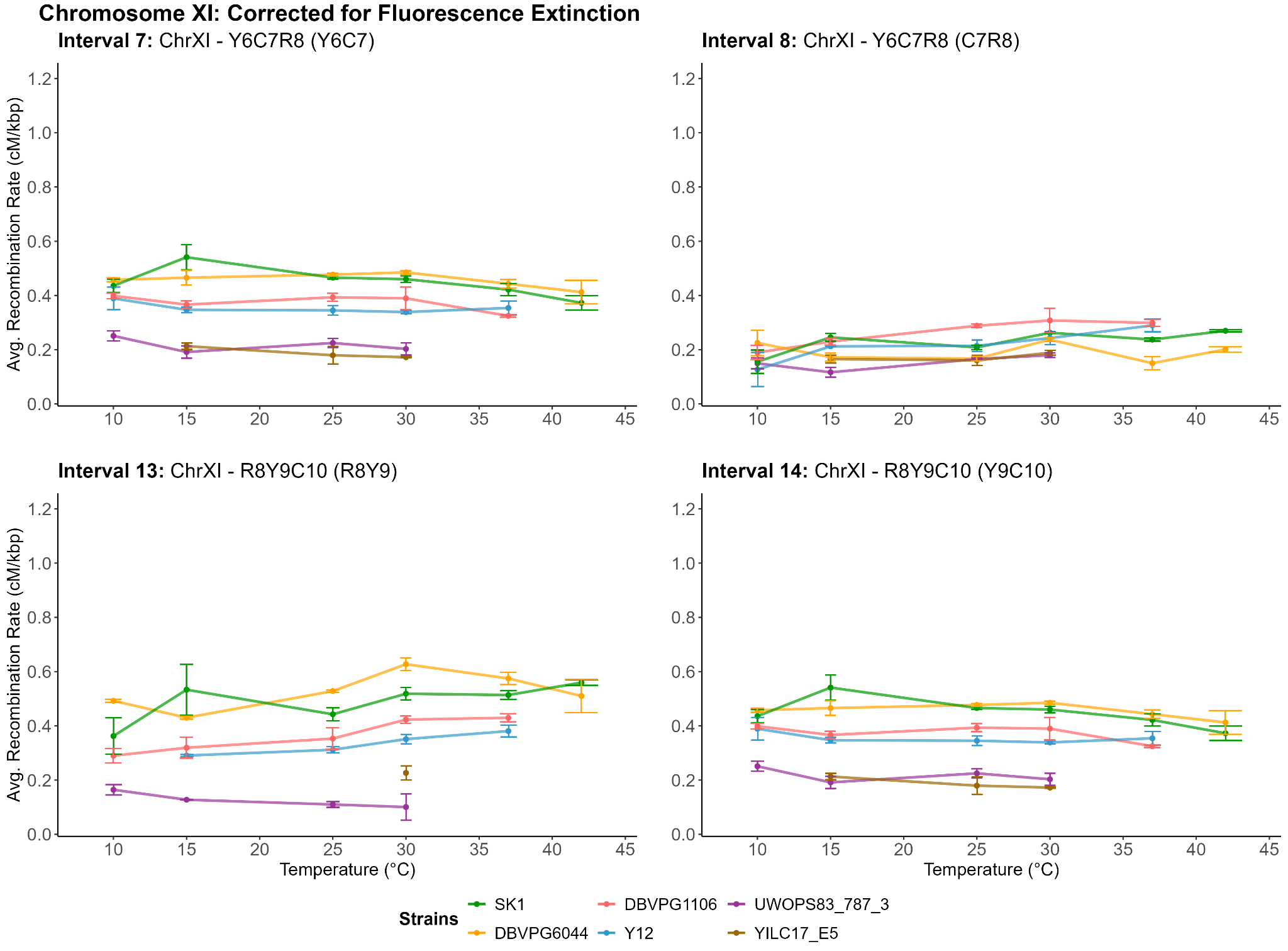

Average recombination rate (in cM/kbp) as calculated for intervals 7, 8 ,13 and 14 from sporulated crosses with the ChrXI-Y6C7R8 fluorescent tester strain (7 and 8) and the ChrXI-R8Y9C10 fluorescent tester strain (13 and 14) at all viable temperatures. Strain name refers to the parent strain crossed with each fluorescent tester to produce a hybrid diploid. Recombination rate estimates were corrected for fluorescence extinction using a maximum likelihood model derived in Raffoux et al 2018a. Error bars indicate standard deviation above and below the mean, as calculated between biological replicates.

**Figure S5**

Average recombination rate (in cM/kbp) as calculated for all measured intervals at all viable temperatures, organized by strain. Strain name refers to the parent strain crossed with each fluorescent tester to produce a hybrid diploid. Recombination rate estimates were corrected for fluorescence extinction using a maximum likelihood model derived in Raffoux et al 2018a. Colors differentiate intervals, while shapes indicate the chromosome on which each interval is located. Error bars indicate standard deviation above and below the mean, as calculated between biological replicates.
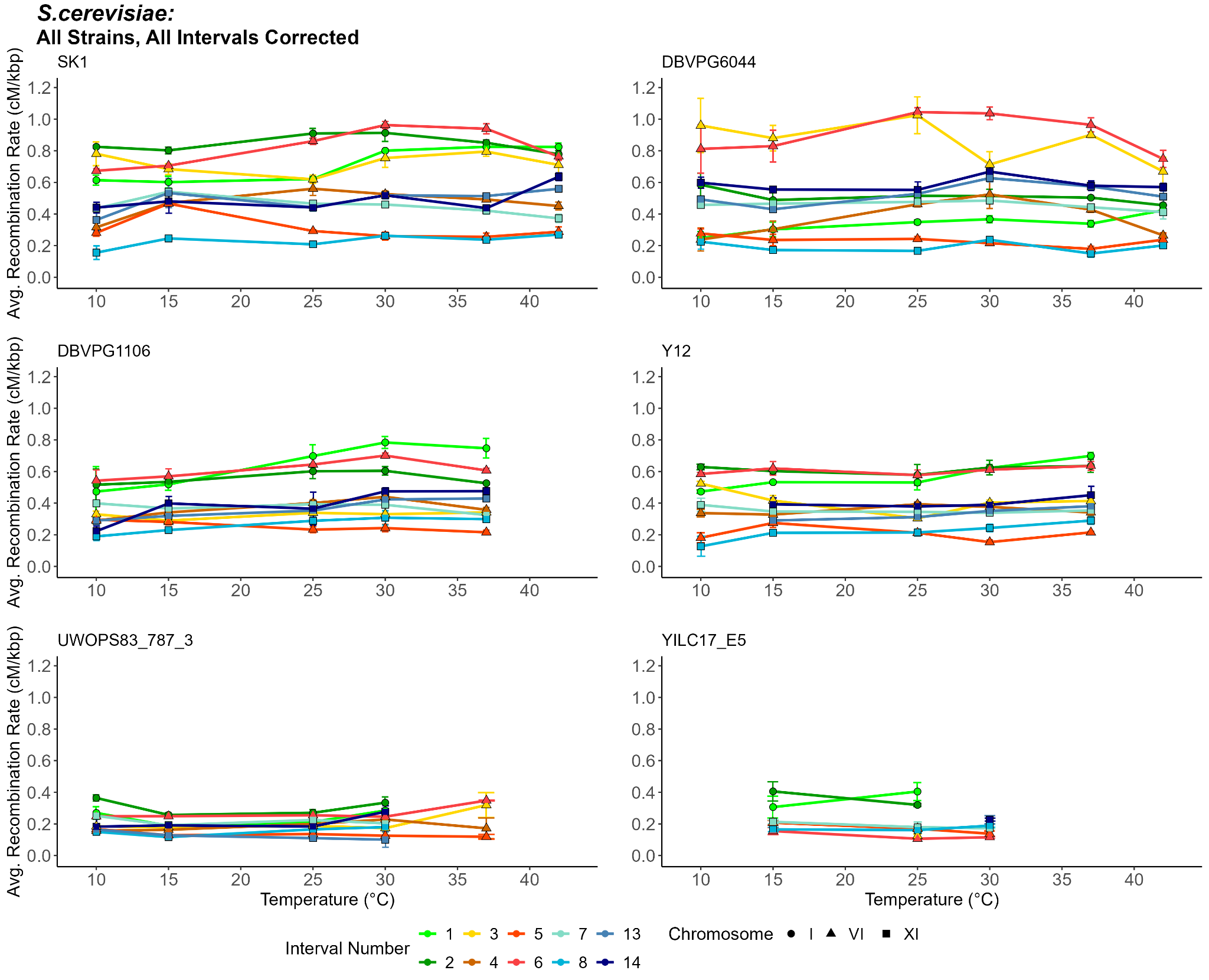

**Figure S6**

Average recombination rate (cM/kbp) as calculated from data pooled across all measured intervals for each *S. cerevisiae* strain at each temperature. In accordance with previous terminology utilized in Raffoux et al 2018b, we refer to this measure as the “global” recombination rate. For our study, this calculation includes a maximum of 10 intervals across chromosomes I, VI, and XI. Note, due to strain-specific complications, not every interval of these 10 was measured at each temperature and included in this calculation.
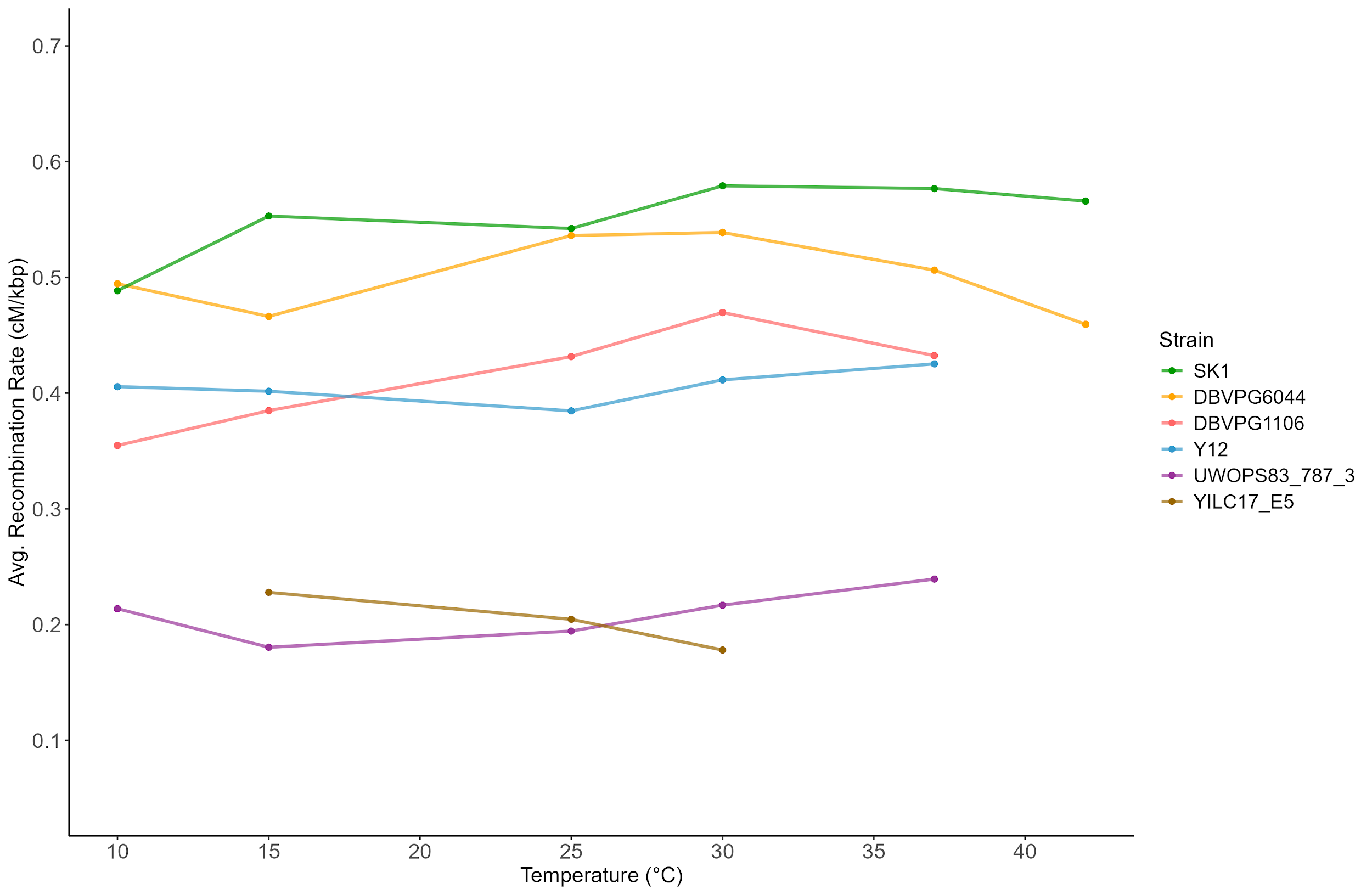

**Text S1**

From Raffoux et al (2018a); maximum-likelihood model that uses exact genotype counts to generate rAB and rBC values corrected for fluorescence extinction in *S. cerevisiae* tester strains with three fluorescent loci per chromosome.

#################################################################

\#	#

\# INRA UMR GQE - Le Moulon RAMDAM group	#

\# Xavier Raffoux, Matthieu Falque	#

\#	#

\#	#

\#	#

#################################################################

###########################################################

\##	F U N C T I O N S

###########################################################

##-----------------------------------------------------------------

```
##  Computes expected frequencies of fluorescence classes for given
##  values of recombination rate in intervals AB and BC,
##  CoC, and fluorescence extinction parameters
```

par2freqs <- function(par) {

rAB <- par[1]

rBC <- par[2]

C <-	par[3]

Aext <- par[4]

Bext <- par[5]

Cext <- par[6]

\#	note that with interference, one has:	(1-rAB)*(1-rBC)	=>	(1-rAB-rBC+C*rAB*rBC)

\#	rAB*(1-rBC)	=>	rAB*(1-C*rBC)

\#	(1-rAB)*rBC	=>	(1-C*rAB)*rBC

\#	rAB*rBC	=>	C*rAB*rBC

fN <-	1/2 * (1-rAB-rBC+C*rAB*rBC) +

1/2 * rAB*(1-C*rBC)*Aext +

1/2 * (1-C*rAB)*rBC*Cext +

1/2 * rAB*C*rBC*Bext +

1/2 * (1-C*rAB)*rBC*Aext*Bext +

1/2 * rAB*(1-C*rBC)*Bext*Cext +

1/2 * rAB*C*rBC*Aext*Cext +

1/2 * (1-rAB-rBC+C*rAB*rBC)*Aext*Bext*Cext

fABC <-	1/2 *(1-rAB-rBC+C*rAB*rBC)*(1-Aext)*(1-Bext)*(1-Cext)

fA <-	1/2 * rAB*(1-C*rBC)*(1-Aext) +

1/2 * (1-C*rAB)*rBC*(1-Aext)*Bext +

1/2 * rAB*C*rBC*(1-Aext)*Cext +

1/2 * (1-rAB-rBC+C*rAB*rBC)*(1-Aext)*Bext*Cext

fB <-	1/2 * rAB*C*rBC*(1-Bext) +

1/2 * (1-C*rAB)*rBC*(1-Bext)*Aext +

1/2 * rAB*(1-C*rBC)*(1-Bext)*Cext +

1/2 * (1-rAB-rBC+C*rAB*rBC)*(1-Bext)*Aext*Cext

fC <-	1/2 * (1-C*rAB)*rBC*(1-Cext) +

1/2 * rAB*(1-C*rBC)*(1-Cext)*Bext +

1/2 * rAB*C*rBC*(1-Cext)*Aext +

1/2 * (1-rAB-rBC+C*rAB*rBC)*(1-Cext)*Aext*Bext

fAB <-	1/2 * (1-C*rAB)*rBC*(1-Aext)*(1-Bext) +

1/2 * (1-rAB-rBC+C*rAB*rBC)*(1-Aext)*(1-Bext)*Cext

fBC <-	1/2 * rAB*(1-C*rBC)*(1-Bext)*(1-Cext) +

1/2 * (1-rAB-rBC+C*rAB*rBC)*(1-Bext)*(1-Cext)*Aext

fAC <-	1/2 * rAB*C*rBC*(1-Aext)*(1-Cext) +

1/2 * (1-rAB-rBC+C*rAB*rBC)*Bext*(1-Aext)*(1-Cext)

freqs <- c(fA, fAB, fABC, fAC, fB, fBC, fC, fN)

names(freqs) <- c("fA", "fAB", "fABC", "fAC", "fB", "fBC", "fC", "fN")

return(freqs)

}

##-----------------------------------------------------------------

\## Computes likelihood of observation given the model with its parameters par

logLike <- function(par, obs) {

if (length(par)==3) par=c(par,Aext=0,Bext=0,Cext=0) #ADD Xext=0 in case of FIT_CONV=FALSE

freqs <- par2freqs(par)

freqs[freqs < 1e-200] <- 1e-200

ll <- sum( obs * log(freqs) )

return(ll)

}

##-----------------------------------------------------------------

\## Finds the set of parameters that maximizes the log-likelihood

fit_params <- function(obs, id, NB_REP_FIT=1000, FIT_CONV=TRUE) {

if (FIT_CONV ) {

lower <- c(rAB = 1e-6, rBC = 1e-6, C = 0.3, Aext = 0, Bext = 0, Cext = 0)

upper <- c(rAB = 0.5, rBC = 0.5, C = 3, Aext = 0.5, Bext = 0.5, Cext = 0.5)

} else {

upper <- c(rAB = 0.5, rBC = 0.5, C = 3)

lower <- c(rAB = 1e-6, rBC = 1e-6, C = 0.3)

}

resReps <- NULL

control <- list(fnscale = -1, factr = 1e3, maxit=100)

methods <- c("Nelder-Mead", "BFGS", "CG", "L-BFGS-B", "SANN")

i1 <- i2 <- i3 <- i4 <- i5 <- i6 <- 1

iRep <- 1

for (iRep in 1:NB_REP_FIT) {

thisInitPar <- rep(NA, length(upper))

for (iParam in 1:length(upper)) thisInitPar[iParam] <- runif(1, lower[iParam], upper[iParam])

res <- optim(par=thisInitPar, lower=lower, upper=upper, fn=logLike, method=methods[4], control=control, obs=obs)

res$par

resReps <- rbind(resReps, res$par)

}

par(mfrow=c(1,length(upper)))

parMode <- rep(NA,length(upper))

for (i in 1:length(upper))	{

dens <- density(resReps[,i])

hist(resReps[,i], proba=T, nclass=NB_REP_FIT/50, main=id, xlab=names(lower)[i], ylim=c(0,max(dens$y)))

lines(dens, col=4)

parMode[i] <- dens$x[which.max(dens$y)]

abline(v=parMode[i], col=2)

}

parMean <- apply(resReps, MARGIN=2, FUN=mean)

parCv <- apply(resReps, MARGIN=2, FUN=function(x) return(sd(x)/mean(x)))

theor <- par2freqs(parMode) * sum(obs)

ssd <- sum((theor-obs)^2)

parCv[is.nan(parCv)] <- 0

return(list(parMean=parMean, parCv=parCv, parMode=parMode, ssd=ssd))

}

###########################################################

\##	M A I N

###########################################################

pdf("Check_Parameters_Fit.pdf", 30,5)

\# obs is a vector with numbers of spores observed for each of the 8

\# classes of fluorescence

obs <- c(

App = 76587,	# A++

ABp = 116232,	# AB+

ABC = 389196,	# ABC

ApC = 16755,	# A+C

pBp = 16191,	# +B+

pBC = 76943,	# +BC

ppC = 116181,	# ++C

ppp = 384696	# +++

)

\# id is sample name (character string) to be displayed in graphs

id <- "MySample"

resFit <- fit_params(obs, id, NB_REP_FIT=1000, FIT_CONV=TRUE)

\# resFit is a list with the following vectors:

\#	parMean (mean of 1000 fits)

\#	parCv (sd/mean of 1000 fits)

\#	parMode (mode of 1000 fits) = best estimate value to use

\#	parMean (mean of 1000 fits)

\#

\# Each vector has 6 elements corresponding respectively to

\# the following 6 parameters :

\#	rAB

\#	rBC

\#	C

\#	Aext

\#	Bext

\#	Cext

results <- resFit$parMode

names(results) <- c("rAB", "rBC", "C", "Aext", "Bext", "Cext",)

print(results)

dev.off()

**Text S2**

Derived in house from Text S1; maximum-likelihood model that uses exact genotype counts to generate rAB values corrected for fluorescence extinction in *S. uvarum* tester strains with only two fluorescent loci per chromosome.

###########################################################

\##	F U N C T I O N S

###########################################################

##-----------------------------------------------------------------

```
##  Computes expected frequencies of fluorescence classes for given
##  values of recombination rate in intervals AB and BC,
##  CoC, and fluorescence extinction parameters
```

par2freqs <- function(par) {

rAB <- par[1]

Aext <- par[2]

Bext <- par[3]

\#	note that with interference, one has:	(1-rAB)*(1-rBC)	=>	(1-rAB-rBC+C*rAB*rBC)

\#	rAB*(1-rBC)	=>	rAB*(1-C*rBC)

\#	(1-rAB)*rBC	=>	(1-C*rAB)*rBC

\#	rAB*rBC	=>	C*rAB*rBC

fN <-	1/2 * (1-rAB) +

1/2 * rAB*Aext +

1/2 * rAB*Bext +

1/2 * (1-rAB)*Aext*Bext

fAB <-	1/2 *(1-rAB)*(1-Aext)*(1-Bext)

fA <-	1/2 * rAB*(1-Aext) +

1/2 * (1-rAB)*(1-Aext)*Bext

fB <-	1/2 * rAB*(1-Bext) +

1/2 * (1-rAB)*(1-Bext)*Aext

freqs <- c(fA, fAB, fB, fN)

names(freqs) <- c("fA", "fAB", "fB", "fN")

return(freqs)

}

##-----------------------------------------------------------------

\## Computes likelihood of observation given the model with its parameters par

logLike <- function(par, obs) {

if (length(par)==1) par=c(par,Aext=0,Bext=0) #ADD Xext=0 in case of FIT_CONV=FALSE

freqs <- par2freqs(par)

freqs[freqs < 1e-200] <- 1e-200

ll <- sum( obs * log(freqs) )

return(ll)

}

##-----------------------------------------------------------------

\## Finds the set of parameters that maximizes the log-likelihood

fit_params <- function(obs, id, NB_REP_FIT=1000, FIT_CONV=TRUE) {

if (FIT_CONV ) {

lower <- c(rAB = 1e-6, Aext = 0, Bext = 0)

upper <- c(rAB = 0.5, Aext = 0.5, Bext = 0.5)

} else {

upper <- c(rAB = 0.5)

lower <- c(rAB = 1e-6)

}

resReps <- NULL

control <- list(fnscale = -1, factr = 1e3, maxit=100)

methods <- c("Nelder-Mead", "BFGS", "CG", "L-BFGS-B", "SANN")

\# i1 <- i2 <- i3 <- i4 <- i5 <- i6 <- 1

\# iRep <- 1

for (iRep in 1:NB_REP_FIT) {

thisInitPar <- rep(NA, length(upper))

for (iParam in 1:length(upper)) thisInitPar[iParam] <- runif(1, lower[iParam], upper[iParam])

res <- optim(par=thisInitPar, lower=lower, upper=upper, fn=logLike, method=methods[4], control=control, obs=obs)

res$par

resReps <- rbind(resReps, res$par)

}

par(mfrow=c(1,length(upper)))

parMode <- rep(NA,length(upper))

for (i in 1:length(upper))	{

dens <- density(resReps[,i])

hist(resReps[,i], proba=T, nclass=NB_REP_FIT/50, main=id, xlab=names(lower)[i], ylim=c(0,max(dens$y)))

lines(dens, col=4)

parMode[i] <- dens$x[which.max(dens$y)]

abline(v=parMode[i], col=2)

}

parMean <- apply(resReps, MARGIN=2, FUN=mean)

parCv <- apply(resReps, MARGIN=2, FUN=function(x) return(sd(x)/mean(x)))

theor <- par2freqs(parMode) * sum(obs)

ssd <- sum((theor-obs)^2)

parCv[is.nan(parCv)] <- 0

return(list(parMean=parMean, parCv=parCv, parMode=parMode, ssd=ssd))

}

###########################################################

\##	M A I N

###########################################################

pdf("Check_Parameters_Fit.pdf", 30,5)

\# obs is a vector with numbers of spores observed for each of the 8

\# classes of fluorescence

obs <- c(

Ap = 2232,	# A+

AB = 3110,	# AB

pB = 2503,	# +B

pp = 3803 # ++

)

\# id is sample name (character string) to be displayed in graphs

id <- "MySample"

resFit <- fit_params(obs, id, NB_REP_FIT=1000, FIT_CONV=TRUE)

\# resFit is a list with the following vectors:

\#	parMean (mean of 1000 fits)

\#	parCv (sd/mean of 1000 fits)

\#	parMode (mode of 1000 fits) = best estimate value to use

\#	parMean (mean of 1000 fits)

\#

\# Each vector has 6 elements corresponding respectively to

\# the following 6 parameters :

\#	rAB

\#	rBC

\#	C

\#	Aext

\#	Bext

\#	Cext

results <- resFit$parMode

names(results) <- c("rAB", "Aext", "Bext")

print(results)

dev.off()
